# Differential transcriptional reprogramming by wild type and lymphoma-associated mutant MYC proteins as B-cells convert to lymphoma-like cells

**DOI:** 10.1101/2020.07.13.200238

**Authors:** Amir Mahani, Gustav Arvidsson, Laia Sadeghi, Alf Grandien, Anthony P. Wright

**Author notes:** The authors contributed equally to this work (Amir Mahani, Gustav Arvidsson). Correspondence: Anthony Wright, Division of Biomolecular and Cellular Medicine, Department of Laboratory Medicine, Novum (level 5), 141 57 Huddinge, Sweden.

## Abstract

The transcription factor MYC regulates the expression of a vast number of genes and is implicated in various human malignancies, for which it’s deregulation by genomic events such as translocation or amplification can be either disease-defining or associated with poor prognosis. In hematological malignancies MYC is frequently subject to missense mutations and one such hot spot where mutations have led to increased protein stability and elevated transformation activity exists within its transactivation domain. Here we present and characterize a model system for studying the effects of gradually increasing MYC levels as B-cells progress to lymphoma-like cells. Inclusion of two frequent lymphoma-associated MYC mutants (T58A and T58I) allowed for discrimination of changes in the MYC regulatory program according to mutation status. Progressive increase in MYC levels significantly altered the transcript levels of 7569 genes and subsets of these were regulated differently in mutant MYC proteins compared to WT MYC or between the mutant MYC proteins. Functional classification of the differentially regulated genes based on expression levels across different MYC levels confirmed previously found MYC regulated functions such as ribosome biogenesis and purine metabolism while other functional groups such as the downregulation of genes involved in B-cell differentiation and chemotaxis were novel. Gene sets that were differently regulated in cells overexpressing mutant MYC proteins contained an over-representation of genes involved in DNA Replication and transition from the G2 phase to mitosis. The cell model presented here mimics changes seen during lymphoma development in the Eμ-Myc mouse model as well as MYC-dependent events associated with poor prognosis in a wide range of human cancer types and therefore constitutes a relevant cell model for *in vitro* mechanistic studies of wild type and mutant MYC proteins in relation to lymphoma development.

## Introduction

The transcription factor MYC plays a central role in the onset and progression of several B-cell lymphomas.[1] The defining characteristic of Burkitt’s lymphoma (BL) is a translocation event where *MYC* is fused to the immunoglobulin heavy chain locus leading to its aberrant expression(t8;14)(q24;q32).[2] Less frequently in BL, *MYC* is translocated to one of two loci encoding for immunoglobulin light chains but the result is nevertheless deregulated expression and a BL phenotype. Aberrant MYC expression is however not sufficient to trigger malignant transformation and secondary events, such as concordant overexpression of Bcl-xL (encoded by BCL2L1) and BMI1 that inhibit the intrinsic apoptotic pathway and the ARF/TP53 axis respectively, are required for MYC-driven B-cell transformation.[3] *MYC* is additionally the most frequently mutated gene in BL.[4] The conserved Myc Homology Box 1 (MB1) within the transactivation domain contains a transiently structured proline-rich sequence where missense mutations frequently occur in BL and other hematological malignancies.[5–9] Among these, mutations most commonly lead to a threonine to isoleucine substitution at residue 58 (T58I), while the somewhat less frequently occurring substitution with alanine at the same residue (T58A) mutation causes higher transforming activity than both wild type (WT) and T58I in cellular model systems.[10] Additionally T58A has been shown to promote B-cell lymphomagenesis in murine models.[10–12] MYC stability is affected by sequential phosphorylation at S62 and T58 where the former activates and stabilizes MYC while a subsequent phosphorylation at T58 primes MYC for proteasome-mediated degradation.[13] Proteins carrying the T58A mutation have been shown to be stabilized while MYC proteins carrying T58I are surprisingly similar to WT in terms of half-life indicating that additional mechanisms affecting MYC protein levels are affected when the threonine residue at position T58 is altered to some other amino acids that cannot be phosphorylated.[5,10]

MYC/MAX heterodimers bind motifs in the regulatory regions of genes with different affinity and thereby control the expression level of genes. While the heterodimer has the highest affinity for canonical E-box motifs (CACGTG) or variants thereof, MYC/MAX have been shown to bind to the regulatory regions of a large number of regulated target genes, including genes with non-canonical E-box motifs or without recognizable E-box motifs in regulatory regions.[14–16] The differing affinity of MYC/MAX for different E-box motifs in different genes allows for specific regulation of target genes in a MYC-level dependent fashion, where the promoters containing high-affinity motifs are preferentially bound at low MYC levels and genes with progressively lower affinity are progressively engaged as MYC levels increase.[16] In a transformed cell with deregulated MYC expression, MYC would be capable of regulating an abnormally large set of genes, ranging from those for which MYC has the highest promoter affinity to those with much lower affinity for MYC.

Previous studies of global gene expression changes upon MYC overexpression have focused on gene expression changes in relation to MYC occupancy in regulatory gene regions[17,18] or in relation to E-box occurrences in regulatory regions upon gradually increasing MYC levels.[16] An *in vivo* study of the Eμ-Myc mouse model of B-cell lymphoma using B-cells at three different stages of lymphomagenesis, each with different MYC levels, identified almost four thousand significantly changed genes between cells representing the different stages.[17] Efforts have also been made to identify genes that are directly regulated by MYC, by combining gene expression profiling with chromatin immunoprecipitation in a B-cell line,[19] or more recently a list of 100 direct MYC targets was identified and subsequently validated across an array of human cancers by employing techniques for rapid induction of MYC degradation in combination with RNA-seq of nascent RNA.[20] Another study investigated global gene expression changes following overexpression of MYC carrying mutations in MBI in relation to WT MYC in rat fibroblasts and found significant differences in gene expression for hundreds of genes between cells expressing WT MYC and the MYC mutants.[21] However, in that study it was difficult to identify functions and pathways differentially regulated by the MYC mutants in relation to WT, possibly due to the cellular context. The study did however describe a differential regulation of Nop56 and a subsequent differential transformation capacity, between cells overexpressing WT MYC and T58I, again suggesting that subtle alterations to the sequence and protein conformation of MYC can have large effects on gene regulation and phenotypic manifestations.

In the present study, we use a transduced primary B-cell model of MYC-driven B-cell lymphoma to identify genes that are regulated as essentially normal murine B-cells transform into lymphoma-like cells in response to progressively increasing MYC levels and further to identify how the lymphoma-associated T58A and T58I mutations affect this transition in terms of altered gene expression programs. Previous studies have shown that analogously modified cells cause lymphomas when injected into mice.[22]

## Results

### Effects of progressively increased wild type and mutant MYC levels on cell growth, proliferation and cell cycle distribution

An *in vitro* cell model for investigating the role of MYC in lymphoma development was developed using LPS-activated primary murine B-cells, transduced with retroviral vectors containing doxycycline-regulated coding sequences for wild type (WT) MYC or one of two lymphoma-associated MYC mutants, in which threonine-58 is substituted with alanine (T58A) or isoleucine (T58I), together with vectors for constitutive expression of the anti-apoptotic proteins Bcl-xL, encoded by *BCL2L1*, and BMI1, as described previously.[3] Concordant expression of the three proteins is necessary for oncogenic transformation of the murine B-cells as they otherwise undergo apoptosis or fail to proliferate. The resulting transformed cells expressed B-cell markers (CD19 and CD45R) but were negative for CD90 (expressed in T-cells but not B-cells), indicating that the transduced cells were predominantly B-cells as expected (Supplementary Figure 1).

To understand changes in cellular characteristics and how they differ between WT and mutants as increasing MYC levels drive the transduced B-cells to Lymphoma-like cells, we overexpressed MYC, T58A and T58I at different levels by progressively increasing the doxycycline concentrations added to the transduced cells for 48, 96 and 144 hours (Figure 1A). The doxycycline-mediated increase in WT and mutant MYC proteins is shown in Figure 1B (see also Supplementary Figures S2 and S8 for a representative image and original data). Consistent with previous studies the T58A mutant reaches somewhat higher levels than the WT and T58I mutant proteins that are expressed at similar levels.[5,10] As shown in Figure 1C, cells can be seen to enter the cell cycle by 48 hours (fewer G_1_/G_0_ cells and more S-phase cells) but there is little or no change in cell growth or proliferation (cell size or number). At later time points (96 h and 144 h) the cells progressively grow (larger size), proliferate (increased number) and enter the cell cycle in response to both MYC level and time. T58A cells outperform WT and T58I cells in these respects by showing greater sensitivity to doxycycline level, presumably due, at least in part, to the higher expression level of the T58A mutant protein. Interestingly, the T58I mutant shows the lowest response with regard to growth, proliferation and cell cycle entry both with regard to MYC level and time.

**Figure 1.**
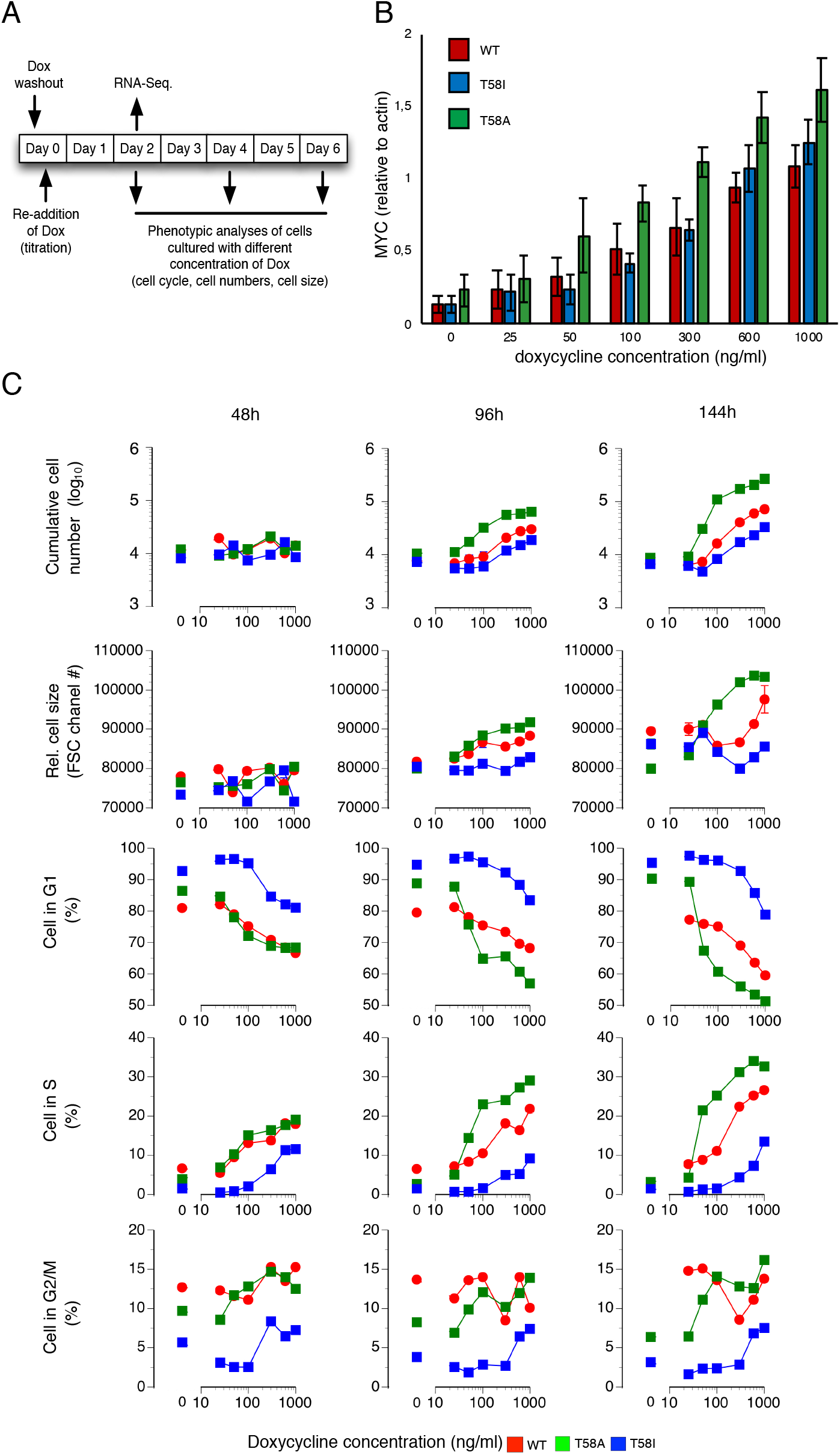
Phenotypic characterization of immortal B-cells expressing doxycycline regulatable MYC. (A) Experimental setup, time points for doxycycline wash out and re-addition as well as time points for data collection and RNA sampling (B) MYC protein levels at different levels of doxycycline in culture medium (0, 25, 50, 100, 300, 600 and 1000 ng/ml) assessed by Western Blot for WT MYC (red), T58A MYC (green) and T58I MYC (blue) in relation to actin levels. (C) Flow cytometry data for WT MYC (red), T58A (green) and T58I (blue) sampled at 48 h, 96 h and 144 h following addition of doxycycline at 0, 25, 50, 100, 300, 600 and 1000 ng/ml. From top to bottom the panels represent: Cumulative live cell number, Relative cell size estimated by the forward scatter channel (FSC) and G0/G1, S and G2/M cell cycle stage assessment for cells stained with propidium iodide (percentages).

### Global gene expression changes in response to progressively increased levels of wild type and mutant MYC proteins

To identify MYC-level associated changes in transcript levels and differences between WT and mutants at a global scale, mRNA was extracted following 48 h of MYC overexpression for seven different levels of MYC expression. While differences in global transcript levels were observed between the B-cells based on *MYC* mutation status, as visualized by the principal component analysis in Figure 2A and B, the largest effect is related to MYC level, where there is a consequent shift to the right along principal component 1 as MYC levels increase while principal components 2 and 3 reveal differences between genotypes. The expression data set was subsequently subject to the glmQLF workflow within the EdgeR package to extract gene models with significant transcript level changes in WT in response to increasing MYC levels as well as for genes that were differentially changed between WT and T58A or T58I. In response to increasing MYC levels, we found 7263 (WT), 347 (WT vs. T58A) and 683 (WT vs. T58I) significant gene models for the respective comparative groups. Selected models were required to have a ≥ 2-fold changed transcript level between at least two conditions (MYC level or mutant status) and an FDR q-value < 0.01 (listed in Supplementary Table S1).

**Figure 2.**
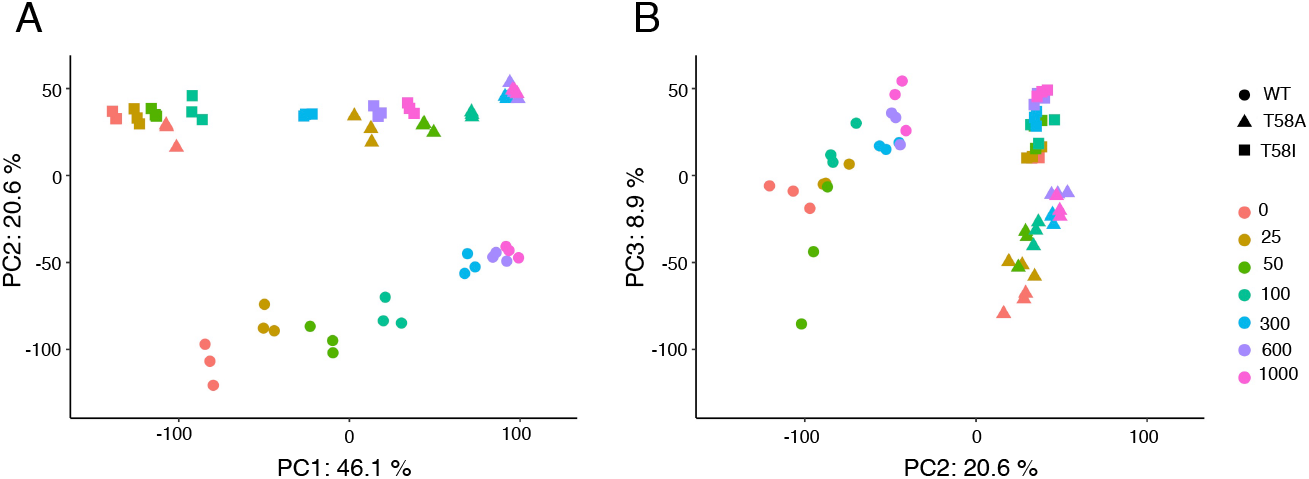
Principal component analysis of global gene expression data. The first three principal components describe 75.6% of the variation in the global gene expression data across the three MYC genotypes and seven expression levels. Levels of MYC WT (circles), MYC T58A (triangles) and MYC T58I (squares) were controlled by titrating in doxycycline at the indicated concentrations (ng/ml, colors). (A) Scatter plot of principal component (PC) 1 versus PC2. (B) Scatter plot of PC2 versus PC3. The proportion (%) of the variation accounted for by each PC is shown.

### Clusters of gene models describe genes with similar patterns of differential gene expression

Agglomerative hierarchical clustering was then used to identify clusters of similar gene models within the combined group of selected gene models, based on mean log_2_ transformed RPKM values for the different MYC levels, for WT and mutant MYC proteins independently (Figure 3). The resulting twelve clusters contained between 55 and 1455 genes and could be divided into two cluster classes depending on whether the overall trend in transcript levels was increasing (clusters c3, c7, c9, c10, c11 and c12) or decreasing (clusters c1, c2, c4, c5, c6, and c8) in response to increasing MYC levels (Supplementary Figure S3). As is evident in Figure 3 some clusters contained genes characterized by high transcript levels (e.g. clusters c5, c7 and c9) while others were characterized by genes with low transcript levels (e.g. clusters c4 and c11). Five clusters were significantly enriched for non-coding genes (c2, c6, c8, c11 and c12) while for six clusters a significant under-representation was observed (c3, c4, c5, c7, c9 and c10). Notably cluster c2, which contained the lowest number of genes and almost exclusively contained non-coding genes, also had significantly shorter genes when compared to non-regulated genes (median feature length: 366 base pairs (bp) compared to 3810 bp). These and other characteristics of the 12 clusters are summarized in Table 1.

**Figure 3.**
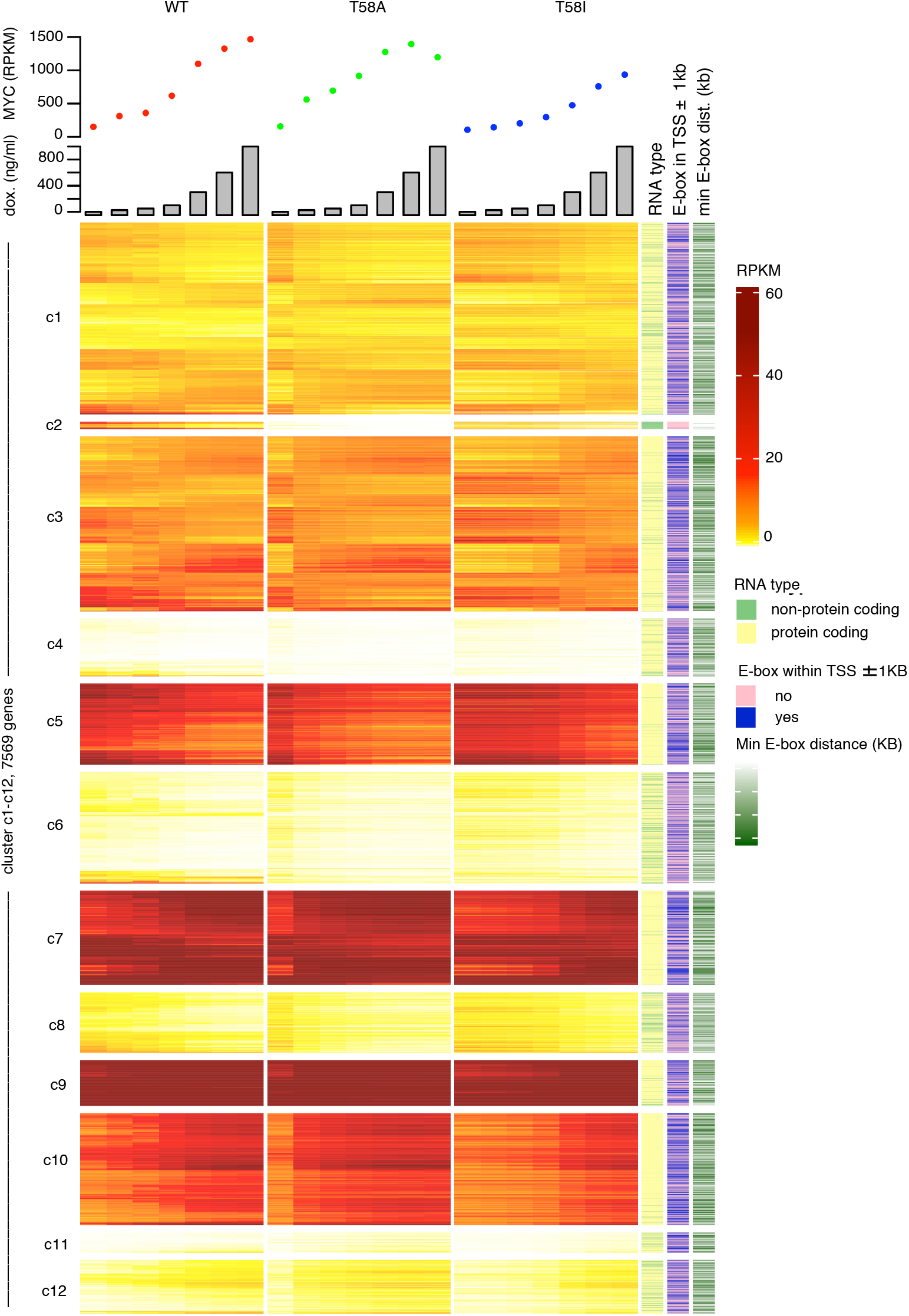
Expression levels for gene clusters Hierarchical clustering based on expression levels for the union of genes that were either significantly changed upon increased MYC levels in WT or the were differentially regulated in response to increasing MYC levels between MYC WT and MYC T58A or between MYC WT and MYC T58I (7569 genes) yielded 12 clusters (n = 3 independent experiments). The heatmap shows reads per kilobase per million of reads (RPKM) for all clustered genes (rows) for different levels (plotted above the heat map) of the three MYC genotypes (MYC WT (red dots), MYC T58A (green) and MYC T58I (blue), columns). Doxycycline concentrations (ng/ml) used to treat the cells are indicated by the bar-chart above the main heatmap. To the right of the main heatmap are three tracks describing, from left to right, whether the differentially regulated genes encode for protein-coding (yellow) or non-coding (light green) RNA, whether (blue) or not (pink) a gene has a canonical E-box motif within 1kb of its transcription start site (TSS) and the last track describes the distance between the TSS and the closest canonical E-box motif.

**Table 1.**
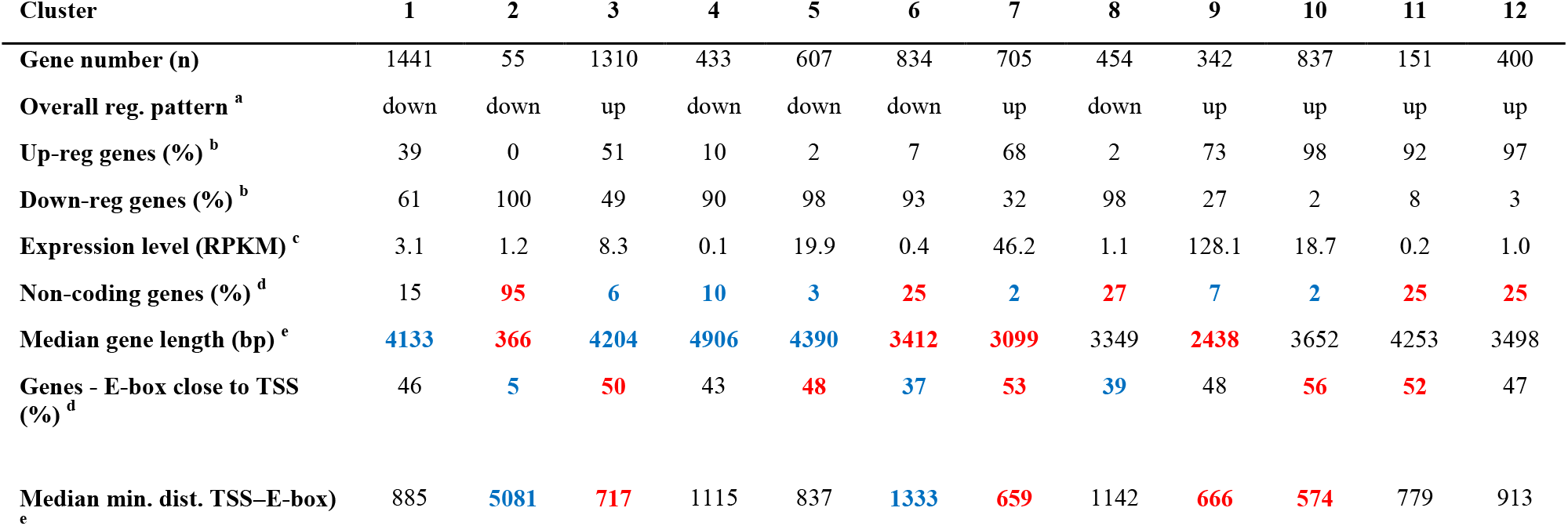
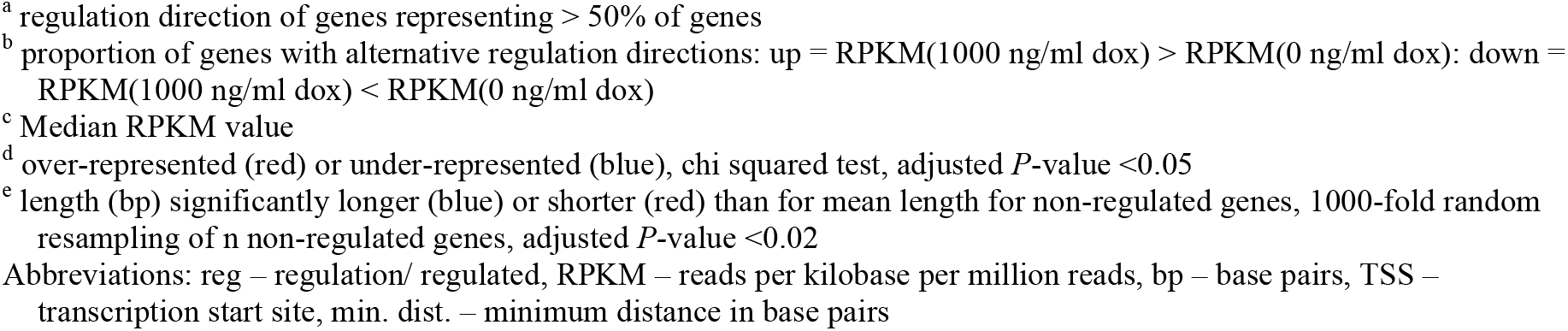
Quantitative characteristics of MYC responsive gene clusters, clustered according to RPKM, MYC response and MYC mutant status.

| Cluster | 1 | 2 | 3 | 4 | 5 | 6 | 7 | 8 | 9 | 10 | 11 | 12 |
| --- | --- | --- | --- | --- | --- | --- | --- | --- | --- | --- | --- | --- |
| Gene number (n) | 1441 | 55 | 1310 | 433 | 607 | 834 | 705 | 454 | 342 | 837 | 151 | 400 |
| Overall reg. pattern <sup>a</sup> | down | down | up | down | down | down | up | down | up | up | up | up |
| Up-reg genes (%) <sup>b</sup> | 39 | 0 | 51 | 10 | 2 | 7 | 68 | 2 | 73 | 98 | 92 | 97 |
| Down-reg genes (%) <sup>b</sup> | 61 | 100 | 49 | 90 | 98 | 93 | 32 | 98 | 27 | 2 | 8 | 3 |
| Expression level (RPKM) <sup>c</sup> | 3.1 | 1.2 | 8.3 | 0.1 | 19.9 | 0.4 | 46.2 | 1.1 | 128.1 | 18.7 | 0.2 | 1.0 |
| Non-coding genes (%) <sup>d</sup> | 15 | 95 | 6 | 10 | 3 | 25 | 2 | 27 | 7 | 2 | 25 | 25 |
| Median gene length (bp) <sup>e</sup> | 4133 | 366 | 4204 | 4906 | 4390 | 3412 | 3099 | 3349 | 2438 | 3652 | 4253 | 3498 |
| Genes - E-box close to TSS (%) <sup>d</sup> | 46 | 5 | 50 | 43 | 48 | 37 | 53 | 39 | 48 | 56 | 52 | 47 |
| Median min. dist. TSS–E-box <sup>e</sup> | 885 | 5081 | 717 | 1115 | 837 | 1333 | 659 | 1142 | 666 | 574 | 779 | 913 |

<sup>a</sup> regulation direction of genes representing > 50% of genes
<sup>b</sup> proportion of genes with alternative regulation directions: up = RPKM(1000 ng/ml dox) > RPKM(0 ng/ml dox): down = RPKM(1000 ng/ml dox) < RPKM(0 ng/ml dox)
<sup>c</sup> Median RPKM value
<sup>d</sup> over-represented (red) or under-represented (blue), chi squared test, adjusted *P*-value <0.05
<sup>e</sup> length (bp) significantly longer (blue) or shorter (red) than for mean length for non-regulated genes, 1000-fold random resampling of *n* non-regulated genes, adjusted *P*-value <0.02
Abbreviations: reg – regulation/ regulated, RPKM – reads per kilobase per million reads, bp – base pairs, TSS – transcription start site, min. dist. – minimum distance in base pairs

Figure 3 also shows differences between the effects of WT and mutant MYC proteins, most notably with regard to T58A which exhibits large changes for many genes, even at the lowest doxycycline concentration where MYC levels are scarcely higher than in WT or T58I. We conclude that the significant gene models describing changes in the data set have the capacity to identify gene expression changes in response to increasing levels of MYC as well as differences in changes between wild type and mutant MYC proteins and between the two mutant MYC proteins.

### Close proximity and higher number of canonical E-boxes near transcription start sites is associated with gene clusters characterized by up-regulated genes

MYC/MAX heterodimers bind with high affinity to canonical E-box motifs (CACGTG) that are generally located closely upstream or downstream of transcription start sites (TSS), leading to activation of transcription.[16] We identified the positions for all canonical E-boxes in the mouse reference genome and derived the distance of each to the nearest TSS. For clusters c3, c7, c10 and c11 there was a significant enrichment of genes with at least one canonical E-box within 1000 bp of the TSS and a significantly shorter median distance between the TSS and the closest canonical E-box compared to non-regulated genes (Table 1). Each of these clusters contain a majority of up-regulated genes in response to increasing MYC levels. Conversely, cluster c2 and c6, characterized by genes with reducing transcript levels in response to MYC level, exhibited the opposite pattern compared to non-regulated genes (i.e. significantly fewer E-boxes ±1000 bp from the TSS and significantly longer median distance between TSS and closest E-box). The enrichment of canonical E-boxes in clusters characterized by genes that are up-regulated in response to increasing MYC levels suggests that in these cells MYC primarily acts as a positive regulator of target genes with high-affinity binding sites.

### Clusters of similar gene models tend to be enriched in genes with distinct functional roles

To functionally classify genes with similar expression profiles in response to increasing levels of WT and mutant MYC, the twelve clusters were subject to enrichment analysis for gene ontology (GO) term gene sets (C5, biological processes) and KEGG pathways (Figure 4, Supplementary Figure S4 as well as Supplementary Tables S2–S3). Enrichment of GO terms were found for all clustered gene sets (FDR adjusted *P*-value ≤ 0.1). Interestingly, there was little overlap between the clusters with regard to the significantly enriched GO terms. In cluster c1 enrichment was primarily found for a select number of down-regulated TOLL-like receptors for lipopeptides and genes with anti-apoptotic functions. Cluster c2 mostly contained non-coding RNA features which resulted in scarce enrichment of GO terms, presumably due to the poorer annotation of non-coding genes and the relatively small size of this cluster. In cluster c3 there was an enrichment of genes implicated in positive regulation of DNA repair and DNA replication as well as gene sets for regulation of the G_2_/M transition of the cell cycle. Cluster c4 genes show enrichment in several GO terms (also enriched in cluster c6) that cannot intuitively be coupled to B-cells but they also show enrichment in the “positive regulation of small molecule metabolic process” term. There is a large number of child terms to this GO term and it is not apparent which of them might be coupled to MYC in this context. Genes associated with the “cytokine–cytokine receptor interaction” KEGG term are also enriched in cluster c4. In cluster c5, enrichment was found for down-regulated B-cell identity genes with functions in B-cell activation and differentiation. The enriched GO terms are not intuitively informative for cluster c6, although notably this cluster is enriched in non-coding RNA transcripts, which are generally poorly annotated. The latter is also true for cluster c8 but here there is also an over-representation of down-regulated genes involved in chemotaxis. Cluster c7, predominantly containing up-regulated genes with high expression values, has an over-representation of genes involved in ribosome biogenesis as well as mitotic nuclear division (also enriched in cluster c10). Cluster c9, principally containing up-regulated genes, has an over-representation of genes associated with purine metabolism while cluster c10 is enriched in genes implicated in non-coding RNA metabolic processes, RNA binding and mRNA transport as well as mitotic nuclear division. Cluster c11, mostly containing up-regulated genes, has a weak enrichment of genes involved in cAMP-mediated signaling. Cluster c12 does not contain genes enriched in processes intuitively related to B-cells but does show weak association to some other GO terms. Notably clusters c11 and c12 contain about 25% non-coding RNAs, including lincRNAs, potential protein-coding transcripts that await validation (TEC) and other transcripts.

**Figure 4.**
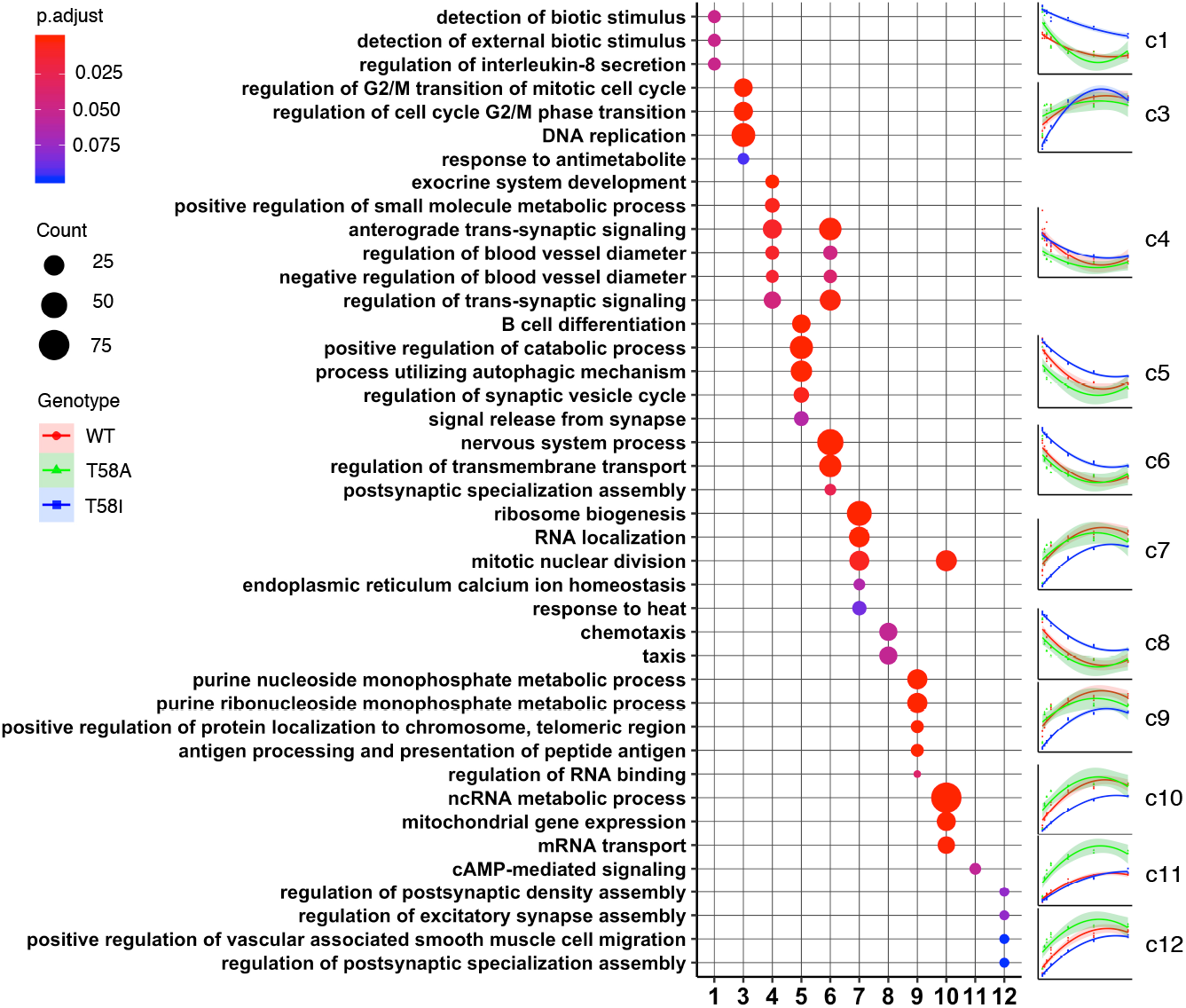
Gene ontology functional classification of clustered genes. Over-representation tests of clustered genes in gene sets with different gene ontology (GO) terms, representing biological processes, for clusters c1 and c3– c12. The most enriched gene sets following GO term simplification are shown. Median RPKM values for genes in denoted clusters are plotted (right panels) for MYC WT (red), MYC T58A (green) and MYC T58I (blue) for increasing doxycycline levels (0, 25, 50, 100, 300, 600 and 1000 ng/ml) the colored lines corresponding to the respective MYC genotype represent a 2nd degree polynomial regression model where the shaded areas indicate the 95% confidence interval for the model. High resolution plots are available in Supplementary Figure S3.

### Comparison of identified genes to independently identified MYC-regulated genes in an in vivo lymphoma model and to direct MYC-target genes

To determine the extent to which the MYC-level associated genes identified in this cell system are related to genes regulated during lymphoma development *in vivo*, we compared our results with existing expression data for the Eμ-Myc mouse model of lymphoma.[17] In that study, Sabo et al. performed RNA-seq analysis (among other analyses) on isolated B-cells at three stages of lymphomagenesis, each with different and progressively increasing MYC levels. After analysis of publicly available data from that study we observed a large overlap (3991 genes, 55%) between the genes with significantly changed transcript levels in response to WT MYC in this study and the genes with significantly altered transcript levels in the study by Sabo et al. (Figure 5A). In addition, the overlap contained 68% of regulated genes identified in the Eμ-Myc model data using the same analysis pipeline as in the present study. In general, the genes regulated during lymphoma development *in vivo* were well represented in the 12 clusters that describe the data from the present study (Figure 5B). Notably, there was a significant over-representation of intersecting genes in clusters c3, c7, c9 and c10, which were functionally classified as containing genes involved in cell cycle, ribosome biogenesis, purine metabolism and pyrimidine metabolism and RNA-related processes (Figure 4).

**Figure 5.**
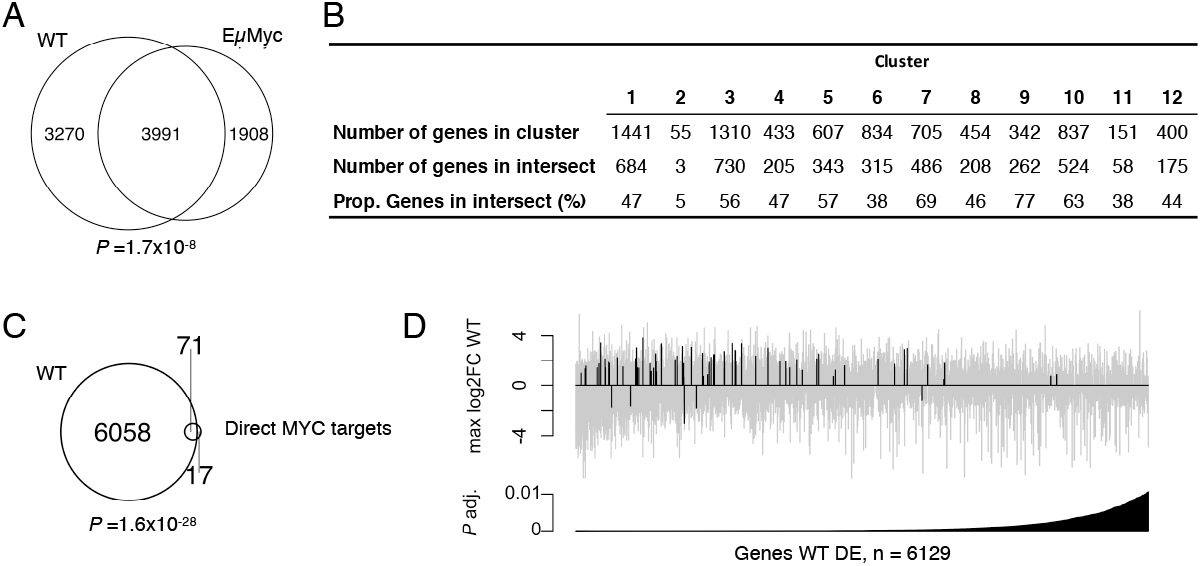
Comparisons with published data sets relevant for MYC-driven lymphogenesis. (A) Overlap between differentially regulated genes in WT MYC and significantly changed genes between any of the conditions measured by Sabó et al. in the Eμ-Myc mouse model.[17] P-value from Fisher’s exact test. (B) Representation of significantly changed genes in the Sabó et al. data set in gene clusters c1–c12. Genes from clusters c3, c7, c9 and c10 were over-represented in the regulated genes from the Sabó et al. study (Fisher’s exact test, P = 0.02, 6,0E-20, 2,0E-20 and 2,0E-09, respectively). (C) Overlap between the 88 of 100 most significant direct MYC target genes derived from Muhar et al.[20] that could be converted to the mouse ortholog with a unique gene identifier and genes with significantly altered transcript levels in WT MYC. P-value from Fisher’s exact test. (D) 6129 genes that were differentially regulated in WT MYC and that had unique gene identifiers following ortholog conversion between human and mouse ranked by adjusted P-value. Each grey bar represents one gene with the 71 genes that were also present in the data from Muhar et al. marked in black. The set of 71 genes was significantly enriched in the set of most significantly regulated genes in WT MYC (Fisher’s exact test, P-value = 0.0012).

Using SLAM-seq to identify direct MYC targets (genes for which the level of nascent transcripts were changed after rapid reduction of MYC levels), Muhar et al. identified a 100 gene signature that correlated well with genes regulated in a wide range of human cancers characterized by highly de-regulated MYC and where high MYC expression is a negative prognostic factor.[20] Eighty-eight of the human signature genes had reliable orthologues in mouse and of these 71 overlapped with the set of significantly regulated genes identified in the present study (Figure 5C). There was also an enrichment of direct MYC target genes among the most significantly changed genes in the present study (Figure 5D). A similar enrichment was found for an intersect of differentially regulated genes from the present study and a list of 668 direct MYC target genes that was identified using a murine B cell line[23] (Supplementary Figure S5).

The 71 genes that were common to the direct MYC target genes identified by Muhar et al. had canonical E-box motifs significantly closer to the TSS than non-regulated genes (median distance: 169 bp as compared to 965 bp for the non-regulated genes, resampling *P*-value = 0.0046). A significant enrichment for the 71 direct target genes was observed in the up-regulated clusters c7, c9 and c10 (Supplementary Figure S6).

We conclude that the MYC-level associated genes identified in this study and particularly the 1886 genes making up clusters c7, c9 and c10, benchmark well in relation to genes that change during lymphoma development in the Eμ-Myc *in vivo* lymphoma model as well as human signature genes that are characteristic for a wide range of human cancers with highly de-regulated MYC expression and high MYC levels as a negative prognostic factor.

### Genes that respond differently in response to MYC level in cells expressing lymphoma-associated MYC mutants or WT MYC

A subset of MYC-level associated genes responded significantly differently to increasing levels of MYC, depending on the mutant status of the expressed MYC protein (WT, T58A or T58I, Figure 6A and Supplementary Table S1). For most genes, the differences in MYC-level associated regulation pattern between cells expressing different MYC proteins do not reach statistical significance (n = 6652 genes, Figure 6A subset A). However, 306 genes show significant changes in relation to MYC level in one or both mutants but not in the WT MYC cells (Figure 6A, subsets B, C and F). A further 611 genes are regulated in cells expressing all three proteins but differentially in one or both mutants compared to WT (Figure 6A, subsets D, E, and G). It is notable that there are many more genes differently regulated in only one of the mutants individually compared to WT (Figure 6A, subsets B, C, D and E, n = 803) than there are genes that are differently regulated in both mutants, relative to WT (Figure 6A, subsets F and G, n = 113).

**Figure 6.**
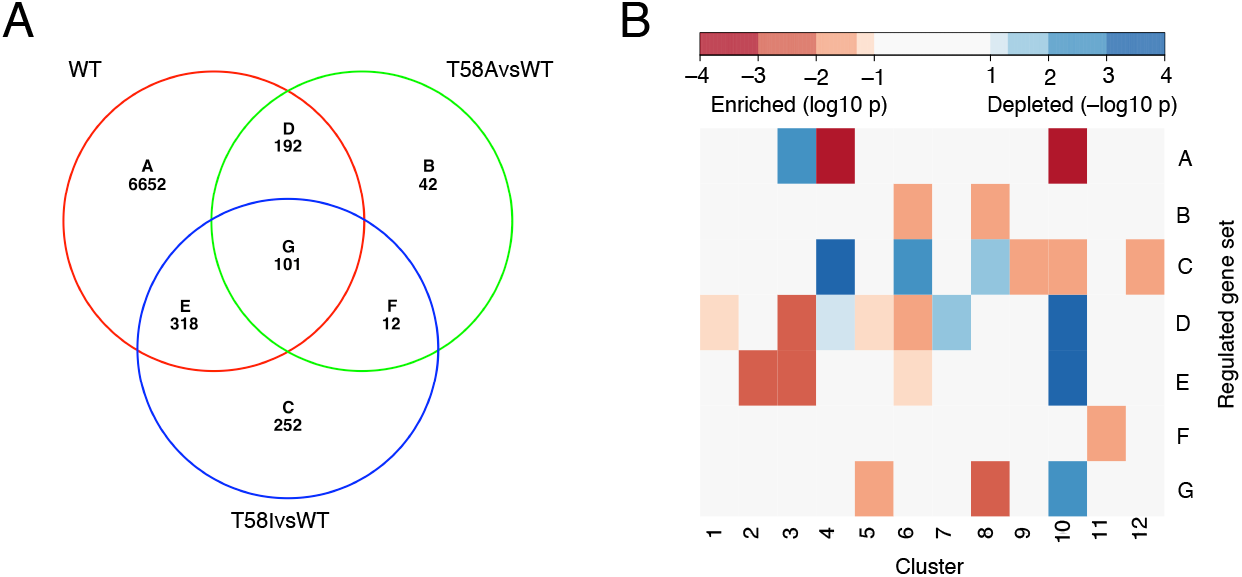
Differences in differentially regulated genes between MYC WT, MYC T58A and MYC T58I. (A) Venn diagram representation of genes with significant transcript level changes (FDR q-value ≤ 0.01 and max fold change ≥ 2). The diagram shows the set of genes with significant regulation in MYC WT (WT) and its overlap with the sets of genes that are regulated significantly differently from WT MYC in MYC T58A (T58A vs. WT) and/ or MYC T58I (T58I vs. WT). (B) Heatmap representation of Fisher’s exact test significance levels for intersects between the Venn diagram subsets in panel A (A–G) and genes in clusters 1–12. Intersects with a significantly larger overlap than expected are represented in red and intersects with an overlap significantly smaller than expected in blue.

An enrichment analysis using the KEGG pathway database for each of the mutant-specific subsets in Figure 6A (subsets B-G) yielded few significantly enriched functional categories of interest, plausibly due to the small size of the gene subsets (Supplementary Figure S7). Instead, subsets A-G were tested for gene intersections with clusters c1-c12 and significant over- and under-representations of genes were found for several intersections in relation to the intersection sizes expected for a random assortment of genes between clusters (Figure 6B). For genes in subset G, where the MYC-level response of genes differed in both mutants relative to WT, there is an under-representation in cluster c10, which otherwise contains a strong over-representation of subset A genes, which are not differentially regulated in cells expressing different MYC proteins. Subset G genes are over-represented in clusters 5 and 8, which are characterized by down-regulated genes involved in e.g. B-cell differentiation and chemotaxis.

The subsets showing most over/under-representation in different clusters are those associated with differences compared to WT for each mutant individually (Figure 6B, subsets B, C, D and E). Interestingly, both mutants individually have genes that are differently regulated compared to WT (subsets D and E) that are over-represented in cluster c3, a cluster which is enriched in genes involved in DNA replication and the entry into mitosis. Interestingly, A-subset genes, which do not exhibit significant differential regulation by the different MYC proteins, are significantly under-represented in cluster c3 genes. Subset D and E genes are instead strongly underrepresented in cluster c10 genes, where subset A genes are strongly over-represented. Taken together this suggests that genes important for mitotic entry and DNA replication tend to be affected by substitution of threonine 58 with alanine or isoleucine, while cluster 10 genes important for e.g. ncRNA metabolic processes, including processing of rRNA and tRNA, tend not to be differentially affected by such amino acid substitutions.

Subset C genes, specifically associated with T58I differences compared to WT, are enriched in several clusters including cluster c9, which is enriched in genes involved in nucleotide metabolism. Moreover, subset C genes are strongly under-represented in cluster c4 genes where subset A genes are strongly over-represented.

In conclusion, there are specific subset-cluster intersections that show significant over/under-representation of intersection genes, showing that there is an underlying non-random structure that describes genes differentially regulated by mutant MYC proteins in relation to WT. These genes tend to be involved in functions such as DNA replication, nucleotide metabolism, mitotic entry, non-coding RNA metabolism, B-cell differentiation and chemotaxis. A second conclusion is that the T58A and T58I mutant MYC proteins appear to be predominantly associated with different, mutation-specific differences in relation to WT.

### Differences in MYC-level associated gene regulation between WT, T58A and T58I

The progressive occupancy of MYC binding sites with progressively reduced affinity in response to increasing levels of MYC has been shown to be important for understanding the normal function of MYC as well as its function in cancer progression.[16] As shown by Figure 3, it is evident that the T58A mutation increases the MYC response sensitivity of many genes. A gene set enrichment analysis (GSEA) was thus performed using a pre-ranked list based on the sensitivity of gene expression to the level of MYC in relation to mutation status (details in materials and methods) together with gene sets from the KEGG pathway database, GO terms for biological processes and a list of curated hallmarks.[24] 29 significantly enriched gene sets were found (adjusted *P*-value ≤ 0.05 and absolute normalized enrichment score ≥ 1.7, Supplementary Table S4). For the majority of the selected gene sets, the sensitivity to the level of MYC was higher for T58A than for T58I and these sets represented different aspects of the cell cycle and genome integrity processes. Only one set (biological process – peptide cross linking) was found where sensitivity was higher in the T58I mutant. A selection of pathways with strong enrichment are presented in Figure 7.

**Figure 7.**
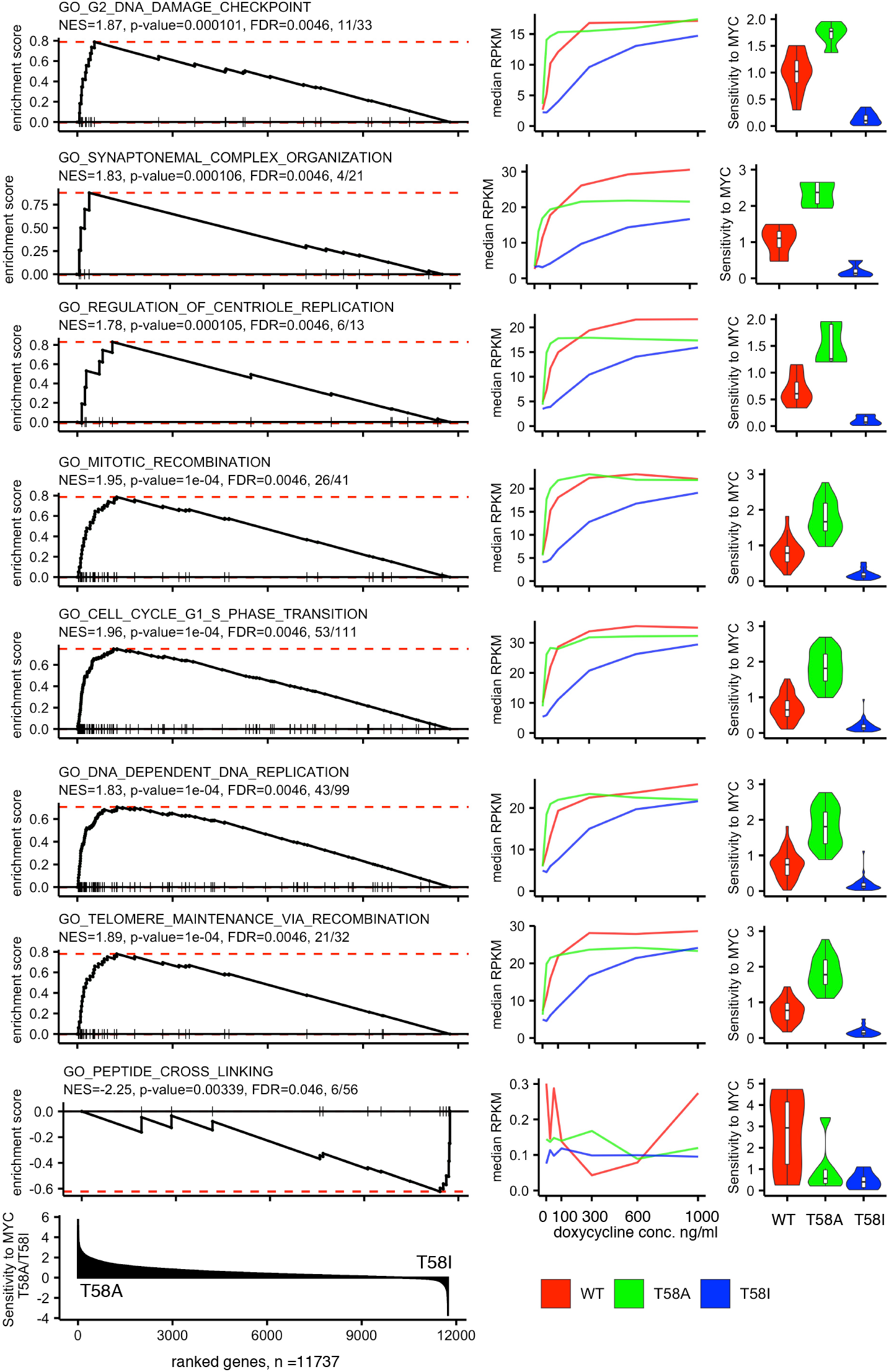
Enriched pathways for genes with sensitivity differences to the levels of MYC T58A and MYC T58I. Representative examples of gene sets that are significantly enriched in a gene set enrichment analysis (GSEA) of genes ranked according to the extent of their sensitivity to T58A MYC in relation to T58I MYC. Left panels: GSEA plots showing a running sum statistic that increases when genes in the ranked list are present in the functional gene set tested. The position of these genes in the rank is indicated by vertical lines. The red dashed line shows the enrichment score value (ES, maximum value of the running sum score). The analysis was performed on 11,737 genes for which human orthologues could reliably be identified and the sensitivity-based rank is shown in the lowest plot. The pathway name is indicated above each plot followed by the normalized enrichment score (NES), GSEA P-value, false discovery rate adjusted P-value (FDR) and the number of genes in the GSEA leading edge (interval between the start of the rank and the gene where ES is attained) in relation to the number of genes in the pathway. Middle panel: Expression-level plots showing median reads per kilobase per millions of reads (RPKM) values for leading edge set of genes identified for each GSEA gene set. Median RPKM values for WT MYC (red), T58A MYC (green) and T58I MYC (blue) at different doxycycline levels (0, 25, 50, 100, 300, 600 and 1000 ng/ml) are plotted. Right panels: Violin plots showing the MYC-level sensitivity of the leading-edge genes in the indicated pathway for WT MYC (red), T58A MYC (green) and T58I MYC (blue). Sensitivity was determined by the change in absolute fold change between 0 to 25 ng/ml and 300 to 600 ng/ml of doxycycline (FC25/0 /FC600/300).

The sensitivity of the T58I mutant protein is lower than WT and T58A with respect to induction of cell cycle entry, cell growth and cell proliferation (Figure 1) and a similar pattern is seen for MYC-dependent regulation of many genes (see Figures 3 and 7). As an alternative way to find genes with specific responses to MYC in T58I, we identified T58I associated genes (subsets C, E, F, G) that differed from WT or T58A by ≥ 1.2-fold at one of the two highest doxycycline concentrations (600 or 1000 ng/ml). Figure 8A shows examples of genes that reflect the predominant pattern in which T58I leads to relatively lower levels of target gene expression. The selected genes represent functions required for cell cycle progression where the lower activity of T58I could contribute to its lower functionality in driving the cell cycle (see Figure 1). However, there are genes showing the opposite pattern in which T58I exhibits a higher level of activity than the other MYC proteins (Figure 8B). The selected group is generally not well annotated but about half the proteins have couplings to protein phosphorylation/ signal transduction and there are functions associated with protein secretion, extracellular matrix and cellular migration. Figure 8C shows that there also genes that show the opposite pattern of MYC-dependent regulation in T58I compared to WT and T58A. Many of the genes are involved in aspects of metabolism, ion homeostasis or transcriptional regulation. For example, the Mnt protein is a competitive inhibitor of MYC due to formation of heterodimers at MYC binding sites with the Max protein. In T58I Mnt is down-regulated, perhaps serving to augment the level of MYC activity, while in WT and to a greater extent in T58A it is up-regulated, perhaps in order to reduce MYC activity. Taken together the results show that the level of response to T58I relative to WT and T58A varies for different classes of genes (Figure 8A–C).

**Figure 8.**
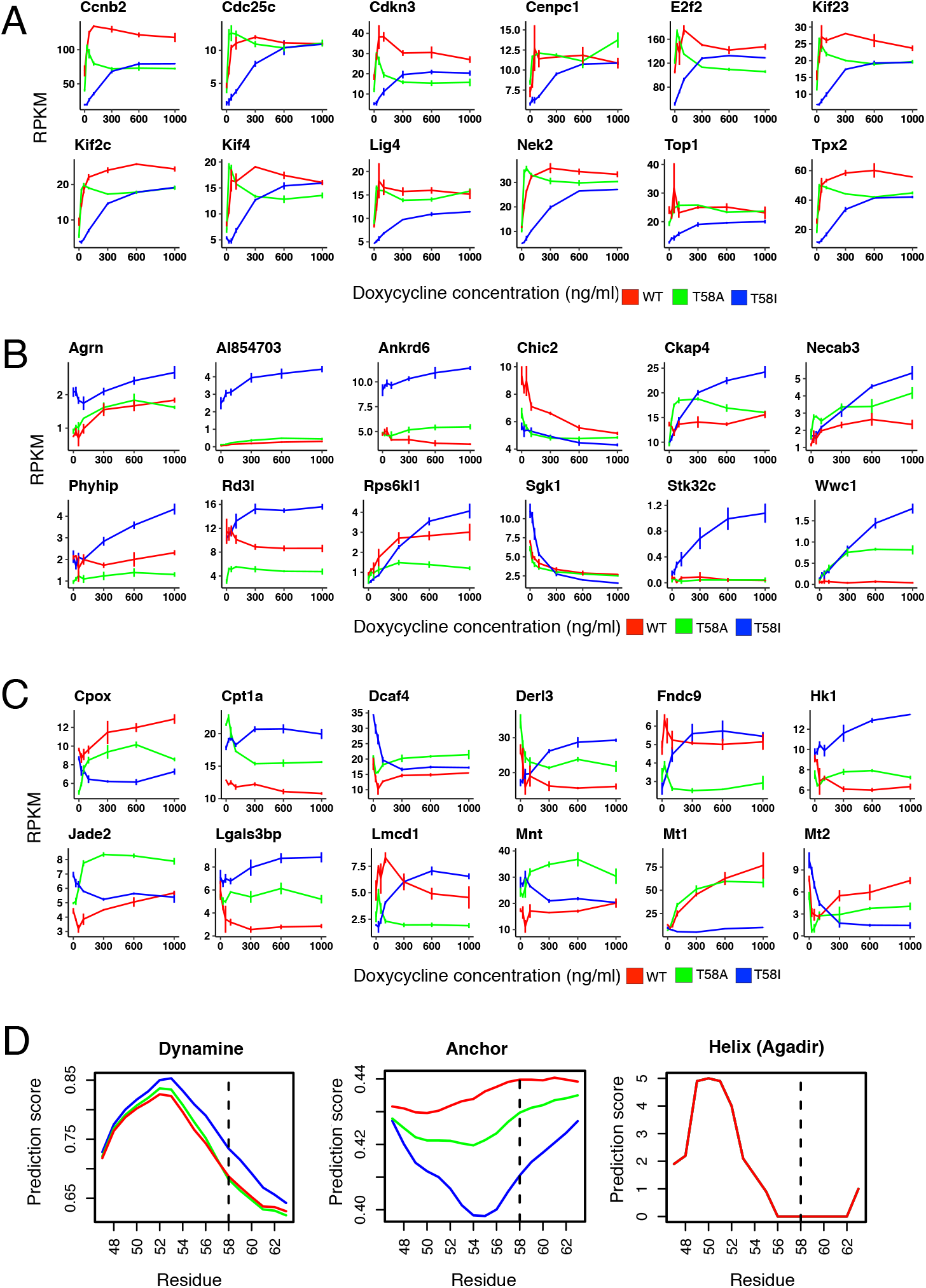
Genes that are differentially regulated by T58I MYC. A) Plots showing the expression levels (RPKM) as a function of doxycycline concentration (0, 25, 50, 100, 300, 600 and 1000 ng/ml) for WT MYC (red), T58A MYC (green) and T58I MYC (blue). (A) Representative genes that show lower regulation by T58I MYC than WT MYC and/ or T58A MYC. (B) Representative genes that show higher regulation by T58I MYC than WT MYC and/ or T58A MYC. (C) Representative genes that show a different regulation direction by T58I MYC than WT MYC and/ or T58A MYC. (D) Protein conformation predictions for the conserved MYC-Box I region, containing residue 58 (dashed line), in WT MYC (red), T58A MYC (green) and T58I MYC (blue). Dynamine predicts rigidity of the protein backbone (scale 0–1 where 1 is highly rigid). Anchor predicts protein interaction propensity (scale 0–1 where 1 is the highest propensity level). Helix (Agadir) is a prediction of alpha-helicity using the Agadir algorithm (the results are essentially identical for the three proteins, explaining the existence of only one visible plotted line).

Different genes require recruitment of different repertoires of co-activator and co-repressor proteins for their regulation and MBI is known to be important for interactions with many partner proteins.[25] Since flexible MYC conformation is important for coupled binding and folding mediated protein interactions,[26,27] it is conceivable that the T58I mutation could affect conformational properties of MBI and thereby its interaction with a subset of MYC partner proteins. Indeed, we showed previously that T58I but not T58A is predicted to affect the level of disordered conformation in MBI.[28] Figure 8D shows that the backbone flexibility (Dynamine) and protein interaction propensity (Anchor) of MBI are predicted to be lower for T58I compared to WT and T58A. There is no effect of the mutations on the predicted alpha-helical propensity of a previously identified transiently alpha-helical region (residues 47–53)[29] immediately N-terminal to residue 58 in MBI (Helix Agadir).

## Discussion

Here we present and characterize roles of MYC in B-cell lymphoma development in B-cells that are primed for lymphoma onset by overexpression of anti-apoptotic factors (BCL2L1 and BMI1) and that therefore progressively adopt a lymphoma-like phenotype as MYC levels are increased. Analogous cells to those described here have been shown to cause lymphoma *in vivo*.[3] The cell system benchmarks well against previous *in vivo* and *in vitro* studies of MYC overexpression and has in addition been used here to better characterize the effects of cancer-associated MYC mutations (T58A and T58I) that are believed to augment the role of MYC in lymphoma development.

Progressive increase in MYC levels was associated with increased cell growth, entry of the transduced cells into the cell cycle and cell proliferation with increased expression of relevant underlying genes. Concordantly, genes related to differentiated B-cell characteristics were down-regulated in response to increasing MYC levels. Notably, gene clusters, defined by changes in transcript levels in response to MYC level and genotype, showed few overlaps in terms of functional categories for which enrichment of genes was found, indicating that data related to expression level, MYC dependence and genotype was sufficient, at least in this case, to identify gene sets with largely different cellular functions. The T58A and T58I mutants were phenotypically different from each other and WT MYC and were associated with largely different subsets of differently regulated genes compared to WT MYC. However, for most genes there was no significant difference in the response pattern for the WT and mutant MYC proteins.

MYC dysregulation is implicated in the onset and progression of several hematological malignancies and its involvement ranges from being disease defining, as it is in Burkitt’s lymphoma,[30] to cases where increased MYC expression, by deregulation or amplification events, is correlated with an inferior clinical outcome.[1] Recent global gene expression studies have largely focused on identifying genes directly regulated by wild type MYC in relation to experimentally detected MYC binding sites or in relation to E-box sequences in P493-6 cells[23,31], U2OS cells[16,18], B- and T-splenocytes[32] or in the Eμ-Myc mouse model[17], collectively showing that MYC has the capacity to increase or decrease the expression of large and diverse, yet discrete sets of genes in different cellular backgrounds. The number of regulated genes upon increased MYC levels in this study (n = 7569) was comparable to previous results using the Eμ-Myc transgenic mouse model and the large overlap with regulated genes in that study indicates that this *in vitro* system faithfully recapitulates major parts of the MYC-driven lymphomagenesis process observed *in vivo*.[17]

Genes regulated in response to MYC overexpression across different model systems have identified common themes related to basal functions such as ribosome biogenesis, RNA processing and biomass accumulation, suggesting a conserved role for MYC in these processes.[33] Indeed the same processes are regulated by the MYC orthologue in *Drosophila melanogaster*.[34] Consistently, MYC has been shown to be a direct regulator of gene transcription by RNA Polymerases I and III, in addition to its role in protein-coding gene transcription, and has thus been suggested to be an overall coordinator of growth-related transcription.[35–37] Mice with either insufficient MYC or reduced ribosome biogenesis have a slower incidence of MYC driven lymphomas [38,39] and an inhibitor of RNA Polymerase I transcription is in clinical trials in patients with hematological malignancies.[40] Consistent with previous studies, clusters c7 and c10 contain genes involved in ribosome biogenesis and rRNA/ tRNA processing with predominantly increasing transcript levels in response to increasing MYC levels.[17,21] Interestingly, promoter regions for genes with functions in ribosome biogenesis also have the highest affinity for MYC binding[16] and in the present study many such genes are highly induced at relatively low MYC levels.

Another study, using RNA-seq measurements of nascent RNA upon abrupt MYC depletion, found that 36% of all factors involved in ribosome biogenesis were directly regulated by MYC as were regulators of adenosine 5′-monophosphate (AMP) metabolism and de novo purine synthesis.[20] These observations are consistent with the present study where an enrichment of genes involved in purine metabolic processes and cAMP-mediated signaling are found in the up-regulated clusters c9 and c11, respectively.[20] Regulation of purine metabolism by MYC has been described before in a B-cell context.^26^ The gene signature with direct MYC targets has been validated across 5583 patient samples representing a diverse set of human malignancies[20] and the observed strong overlap with the present study serves as another strong validation for this model system and the results obtained in relation to lymphomagenesis.

The observation that canonical E-boxes in the predominantly up-regulated clusters c3, c7, c9, c10 and c11 were more frequent close to the TSS of genes or were closer to the transcription start sites than for non-regulated genes, was indicative of a likely direct regulation of many of these genes by MYC. This conclusion is further supported by the significant enrichment of the experimentally determined direct target genes[20] in clusters c7, c9 and c10. Cluster c5 is also enriched in genes with canonical E-boxes close to the TSS but almost all genes in this cluster are down-regulated. Low affinity E-boxes have been associated with genes that are down-regulated in response to activated MYC[16] but the potential role of MYC as a direct repressor of gene regulation remains to be conclusively established. The *P*-value for cluster c5 is close to the significance threshold and thus our data do not contribute strongly to resolution of this issue.

Similar to a previous report using fibroblast-derived cell lines[21] we observe that MYC mutated at residue T58 regulates the majority of genes similarly to WT MYC while a significant subset of genes are differentially regulated in response to increasing levels of the T58A and T58I mutants compared to WT MYC. Interestingly, the differentially regulated gene subsets were largely different for each MYC mutant and thus the effect of substituting T58 cannot be fully attributed to loss of phosphorylation at this site. Our results show that different T58 substitution residues have different effects on the transcriptional activity of the T58A and T58I mutants in this B-cell lymphoma model and this is likely to be coupled to their different phenotypes, consistent with previous studies in other contexts.[10,12,21]

The T58A mutant enhances MYC-dependent induction of cell growth, proliferation and cell cycle entry and this is likely linked to the many genes that tend to be more sensitive to T58A such that lower levels of doxycycline are required for their regulation, compared to WT or T58I MYC. Interestingly, gene Cluster c3 that is characterized by cell-cycle related functions, is over-represented in a set of genes that are differently regulated in T58A compared to WT and T58I MYC expressing cells, while Cluster c10, characterized by cell growth related functions, is depleted in these genes. Thus, we have for the first time been able to identify sets of differentially regulated genes with functions that can account for phenotypic differences that distinguish T58A expressing cells from cells expressing WT or T58I MYC proteins. The T58A mutation has previously been reported to uncouple the role of MYC as an inducer of cell growth and proliferation from its induction of apoptosis during lymphoma onset[12] but in the present study apoptosis is inhibited in all cells by over-expression of anti-apoptotic proteins. Thus, in this lymphoma model increased promotion of cell growth and proliferation, probably resulting from enhanced MYC-induced cell cycle entry, seems to be the best explanation for how T58A contributes to lymphoma development.

Previous studies have shown different and apparently counter-intuitive phenotypic effects of T58I compared to T58A.[10,21] For example, T58I is the most common BL-associated mutation and T58A is rare but T58A is associated with higher tumorigenic activity than T58I. One suggested explanation was that these previous studies were performed in fibroblast-derived cell lines, while BL is a disease of B-cell origin. A major conclusion of this work is that T58I MYC is generally a less active regulator of genes than WT MYC, while T58A MYC is more active, differences that were also seen at the phenotypic level. Thus, this study in a MYC-dependent lymphoma model confirms the differential effects of the T58A and T58I mutations. The apparent reduced activity of T58I may be associated with the reduced phosphorylation of S62 that is observed in this mutant but not T58A, since S62 phosphorylation is required for activation of MYC.[10]

It is important to note that in lymphoma, mutations of T58 are seen in the context of MYC proteins that are overexpressed due to genetic rearrangements. Since MYC induces processes that are both positive (e.g. cell cycle entry, cell growth and cell proliferation) and negative (e.g. apoptosis) for tumorigenesis, one strategy for optimal tumorigenesis maybe to establish a level of MYC activity that optimizes the balance between positive and negative processes. Indeed, there is evidence that positive process genes are activated by lower levels of MYC than negative process genes.[42] In a scenario where MYC is over-expressed at higher than optimal levels, there would be an opportunity for mutations that reduce MYC activity, thus optimizing its tumorigenicity. This would thus be one way to account for the common occurrence of reduced-activity mutations like T58I in lymphomas, particularly those like BL where MYC activation is known to be the primary driver.

A second possible explanation for the role of T58I in tumorigenesis, which is not mutually exclusive to the first, is that T58I enhances MYC-dependent activation of important tumorigenic processes, other than cell growth and proliferation, by activating a subset of genes more efficiently than WT MYC. For example, migration towards and adherence to stromal cells in microenvironments has been shown to enhance the survival properties of some lymphoma cells (see [43] and references therein). While T58I is less active than WT and T58A MYC for regulation of most genes, the group of genes that are differentially regulated in T58I compared to WT does contain a set of genes that are more efficiently regulated by T58I or where the direction of regulation differs in T58I compared to WT or T58A MYC. We were not able to show statistical support for over-representation of genes with particular functions in this gene set but the set contains genes involved in signaling and gene regulation processes as well as metabolism, ion balance, cell migration and cell adhesion.

It was suggested previously that the T58A and T58I mutations may cause different changes to local protein conformation, leading to quantitative or qualitative changes in the interaction of MYC with partner proteins.[10] Consistently there is evidence that the MYC N-terminus is a conformationally disordered region and that conformational changes accompany interaction with partner proteins as part of a coupled binding and folding interaction mechanism.[9,26,44,45] Recently, the stability of protein regions with transient alpha-helical conformation has been shown to be an important modulator of activity for this kind of protein domain, see [46] and references therein, but neither T58A nor T58I is predicted to affect the transient alpha-helical region immediately N-terminal of T58. However, the Dynamine predictor of protein backbone dynamics predicts an increased backbone rigidity in the T58I mutant that is not seen for T58A and the ANCHOR predictor predicts a greater reduction in protein interaction propensity for T58I than for T58A. It is thus possible that the transient extended proline-rich structure reported between residues 55-63 previously,[9] which is characterized by restrained backbone dynamics, is stabilized in T58I but not T58A, leading to a more rigid conformation with lower propensity for coupled binding and folding. This provides a hypothetical model for differential effects of isoleucine and alanine substitutions of T58 on qualitative or quantitative aspects of interactions between MYC and partner proteins.

In summary, we have shown phenotypic and gene regulatory effects of increasing MYC expression as B-cells transit into lymphoma cells in a MYC-dependent manner. We have further shown that different mutations affecting T58 use different strategies to augment MYC-mediated tumorigenesis and that these differences may result from differential effects of the mutations on a transient structure element located in the intrinsically disordered activation domain in the MYC N-terminus. The differential effects potentially cause different effects on the interaction between MYC and partner proteins involved in target gene regulation.

## Materials and Methods

### Preparation of mouse primary B-cells

Ethical permission for the use of mouse primary B-cells was obtained from the Stockholms norra djurförsöksetiska nämnd (Dnr. N375/12). Following local and national guidelines, splenic cells were prepared and stimulated with 25 μg/ml of LPS (Sigma-Aldrich) in RPMI medium with supplements, as described previously.[3]

### Retrovirus vector construction

Vectors containing human wild type *MYC*, *T58AMYC* and *T58IMYC* were generously provided by M.D Cole (Department of Genetics, Geisel School of Medicine, Dartmouth, USA). Human WT MYC, T58A and T58I sequences were amplified by PCR using SK primers containing *MLuI* digestion sites and cloned into *MLuI*-cleaved pSIR-TRE-IRES-EGFP-PGK1-rtTA2 retrovirus expression vector (generously provided by Kari Högstrand), generating pSIR-TRE-MYC-IRES-EGFP-PGK1-rtTA2 vectors. The constructs were confirmed by DNA sequencing. The *BCL2L1* and *BMI1* retroviral expression vectors have been described previously.[3] In the present study, the vector containing *BCL2L1* (encoding for Bcl-xL) encoded EGFP in place of DsRed-Monomer.

### Retroviral transduction

Phoenix-Eco packaging cells (kindly provided by GP. Nolan, Stanford University, maintained in Dulbecco’s modified Eagle’s Medium (DMEM) with 10% FBS at 37°C and 5% CO_2_) were transiently transfected using Lipofectamine 2000 (Life Technologies). Retroviral particles were obtained and concentrated by centrifugation of the conditioned media (6000 × g overnight at +4°C). LPS stimulated B-cells were transduced by retroviral pool spin infection in 8 μg/μl of polybrene (Sigma-Aldrich). Each retroviral pool contained vectors expressing, *WT MYC*, *T58A MYC* or *T58I MYC* as well as both *BCL2L1* and *BMI1*. Cells were re-plated in complete culture medium supplemented with 2 μg/ml doxycycline.

### Cell culture and dose dependent expression of MYC

All three transduced cell lines (WT MYC, T58A MYC and T58I MYC) where cultured in GlutaMAX-RPMI (Gibco) supplemented with 10% FBS (0.01 M HEPES, 1x sodium pyruvate, 0.05 mM 2-beta-mercaptoethanol (Gibco) and 0.1 mg/ml Penicillin-streptomycin (Sigma) with addition of 2 μg/ml doxycycline. For titration experiments, each cell line was removed from doxycycline medium, washed with PBS(3x) and RPMI (3x), and subsequently cultured under 7 different concentrations of doxycycline (0, 25, 50, 100, 300, 600 and 1000 ng/ml) for 48, 96 or 144 hours at 37°C and 5% CO_2_.

### Flow cytometry

Cells expressing WT MYC, T58A MYC or T58I MYC were stained for CD19, CD45R and CD90 markers using monoclonal antibodies, as described previously.[3] Cell cycle analysis was performed with propidium iodide staining, as described previously.[47] Relative cell size was quantified based on the mean fluorescence intensity of the forward scatter. Cell marker and cell cycle samples were analyzed using a BD LSRFortessa Cell Analyzer (BD Biosciences), and acquired data was passed to FlowJo software (v10.xx, Tree Star, Inc.) for data analysis.

### Western Blot

10^6^ cells were collected and directly lysed using 4× LDS sample buffer (Thermo Fisher Scientific) prior to loading onto SDS-PAGE (4–12% Tris-Glycine) gels. Western Blot reagents, including gels and Nitrocellulose membranes for the iBlott Dry-blotting system were purchased from Life Technologies. Anti-Actin (1:100,000, Sigma) and Anti c-MYC (1:1000, 9E10, Thermo Fisher Scientific) antibodies were used to detect proteins.

### RNA isolation, library preparation and RNA-sequencing

Total RNA was isolated using the RNeasy kit (Qiagen) and libraries for NGS were prepared using the TruSeq2.0 sample preparation kit (Illumina) according to manufacturer’s instructions and included a poly-A selection step using poly-T oligo attached magnetic beads. The libraries were thereafter multiplexed on an Illumina HiSeq2000 instrument generating on average 28.6 million 50 bp single-end reads per sample.

### RNA-seq alignment

Read quality was assessed by FastQC (v.0.10.1). The mouse reference genome, build GRCm38.87, was obtained from the ensemble repository with the corresponding gtf annotation file. The reference genome was concatenated with genomic sequences for the protein coding sequences of the four genes present in the retroviral vectors used in the study; human *MYC*, *BCL2L1* and *BMI1* as well as *EGFP* and subsequently indexed using STAR (v.2.5.1b). The 50 bp reads were then aligned to the mouse genome using the splice-aware short read aligner STAR with default options and --outFilterMultimapNmax set to 1 and --sjdbGTFfile pointing to the GRCm38.87 gtf file appended with lines corresponding to the four added features.[48]

### Gene expression and gene cluster profile analyses

Reads per feature (concatenated exons for one gene) were obtained using the featurecounts function from the subread package (v1.5.2) with default options and -g set to “gene_id” and -t set to “exon”.[49] The count table was imported into the R environment (v.3.5.0) and genes with counts per million ≥ 0.95 for at least 3 samples were included in the analysis (n = 14142). Tests for differential expression were conducted using the Bioconductor (v3.8) package edgeR (v.3.24.3) using the glmQLF framework.[50] The differentially expressed features were subsequently subject to hierarchical clustering (Euclidian distances and Ward’s method) using mean log_2_ transformed values for each dose and genotype. Functional classification of the clustered features was conducted by enrichment tests within the databases for GO Biological Processes and KEGG pathways using clusterProfiler (v.3.10.1).[51]

### Distances to canonical E-boxes

The mouse reference genome GRCm38.87 was indexed for Bowtie1 (v1.2.0)[52] followed by alignment of the canonical E-box 6-mer motif (CACGTG) with default options plus the -a flag and -v set to 0, yielding 536,520 uniquely aligning motifs. After conversion of the output file via .bam to .bed using samtools (v1.6)[53] and BEDtools (v2.27.1, https://bedtools.readthedocs.io) respectively, distances to closest known feature for each E-box location were derived using the annotatepeaks.pl script from the Homer package for features in the GRCm38.87.gtf file (http://homer.ucsd.edu/homer/ngs/annotation.html).

### Overlap with published datasets

Supplementary Table 1 from Sabó et al. (Nature 2014)[17] was downloaded and filtered for genes with significant transcript level changes between any of the conditions (FDR q-value ≤ 0.05). The resulting list of genes was subsequently used to look for significant overlaps with all the significantly changed genes for WT MYC from the present study as well as overlaps with each of the twelve clusters. Fisher’s exact tests were used to assess the significance of overlaps.

A list containing the 100 most significantly regulated genes upon attenuation of MYC expression from Muhar et al. (Science 2018)[20] was downloaded from Supplementary Table S3. The list of human genes was subject to orthology conversion using the biomaRt Bioconductor package. 88 of the 100 genes could be converted to a unique mouse identifier and these were subsequently used for comparisons with the present study. A similar approach was taken for a set of 668 direct MYC target genes published by Zeller et al. (PNAS 2006)[19] and presented in Supplementary Figure S5.

### Gene set enrichment analysis

A ranked list of all genes included in the analysis based on their sensitivity to MYC level in relation to mutational status was generated by taking the ratio of change in fold change (FC_dox 25/0_/FC_dox 600/300_) between the mutants (T58A/T58I). The pre-ranked list was used for gene set enrichment analysis using the R package fGSEA(v1.8.0) with gene sets downloaded from the MSigDb (http://software.broadinstitute.org/gsea/msigdb/index.jsp) for the KEGG[54] pathway database (c2.cp.kegg.v6.2.symbols.gmt), gene ontology biological processes (c5.bp.v6.2.symbols.gmt) and hallmark hallmark gene sets [24].

### Protein conformation prediction

Protein conformation characteristics were predicted using web-based prediction software as follows. Alpha-helical propensity was predicted using the Agadir predictor (http://agadir.crg.es).[55] Protein backbone dynamics were predicted using the Dynamine predictor (http://dynamine.ibsquare.be).[56] Protein interaction propensity was predicted using the ANCHOR predictor (https://iupred2a.elte.hu).[57]

## Author Contributions

AM and APW conceived the study with contribution from AG. AM and APW designed the study with contribution from AG and GA. AM generated the MYC-overexpressing cell lines, conducted phenotype analysis and extracted RNA for next generation sequencing with contributions from LS and AG. GA analysed the gene expression data with contributions from APW and AM. GA, AM and APW wrote the manuscript. All authors edited the manuscript and approved of the final version.

## Acknowledgments

The study was supported via grants from: The Swedish Research Council, (APW), The Swedish Cancer Society (APW) and Karolinska Institutet (APW). Processor-intensive computations were performed on resources provided by the Swedish National Infrastructure for Computing (SNIC) at UPPMAX. We would also like to thank the core facility at Novum, BEA, Bioinformatics and Expression Analysis that generated the NGS libraries and preformed the sequencing. It is supported by the board of research at the Karolinska Institute and the research committee at the Karolinska hospital.

